# Guanylate-Binding Proteins Promote Host Defense Against *Leishmania major* by Balancing iNOS/Arg-1 in Myeloid Cells

**DOI:** 10.1101/2025.06.26.661809

**Authors:** Lucy Fry, Het Adhvaryu, Hayden Roys, Anne Bowlin, Gopinath Venugopal, Jordan T. Bird, Charity Washam, Masahiro Yamamoto, Jörn Coers, Stephanie Byrum, Daniel Voth, Tiffany Weinkopff

**Affiliations:** Department of Microbiology and Immunology, College of Medicine, University of Arkansas for Medical Sciences, Little Rock, AR, USA 72205; Department of Biochemistry and Molecular Biology, College of Medicine, University of Arkansas for Medical Sciences, Little Rock, AR, USA 72205; Arkansas Children’s Research Institute, Little Rock, AR, USA 72202; Department of Immunoparasitology, Research Institute for Microbial Diseases, Osaka University, Yamadaoka, Suita, Osaka, Japan; Departments of Molecular Genetics and Microbiology and Integrative Immunobiology, Duke University Medical Center, Durham, NC, USA, 27710

**Keywords:** Leishmaniasis, *Leishmania*, Guanylate binding proteins, Macrophages

## Abstract

Cutaneous leishmaniasis (CL) is a debilitating neglected tropical disease characterized by lesions that can range from self-healing to permanent disfigurations. A predominant Th1 response, which stimulates IFN-γ production, is crucial for parasite control during self-healing CL. While IFN-γ primarily activates macrophages to produce nitric oxide via inducible nitric oxide synthase (iNOS) leading to parasite control, IFN-γ also activates other downstream pathways involved in cell autonomous immunity. One such pathway is the activation of guanylate binding proteins (GBPs), a class of interferon inducible GTPases. However, the role of GBPs during CL has been minimally explored. Utilizing RNA-Seq we found that *Leishmania major* infection leads to the upregulation of several GBPs in C57Bl/6 mice. In vitro studies using GBPChr3 knockout (KO), and C57Bl/6 control mice reveal that bone marrow-derived macrophages (BMDMs) from KO mice exhibit higher parasite burdens following IFN-γ treatment, independent of GBP localization to the parasite. Single-cell RNA-Seq identifies macrophages as the primary expressers of GBPs during *L. major* infection *in vivo*. *In vivo*, GBPChr3 KO mice display increased disease severity and parasite load. GBPChr3 KO macrophages and monocytes show elevated ARG-1 and reduced iNOS expression, indicating a shift toward a parasite-permissive environment that supports parasite growth. These findings highlight a critical role for GBPs in immune-mediated control of CL.

## Introduction

Cutaneous leishmaniasis (CL) represents the spectrum of diseases caused by vector-transmitted *Leishmania* protozoan parasites endemic to the tropical and subtropical regions of the world (1). Each year there are 1-2 million new cases of CL with 12 million ongoing infections in areas including Central and South America, the mediterranean basin, the middle East, and central Asia (2, 3). The species of parasite and the host immune response dictates clinical disease within the spectrum of CL. Both overactive and insufficient immune responses can lead to chronic disease (4). Specifically, control of the intracellular infection is dependent on an effective CD4^+^ Th1 immune response (5). Inability to initiate a dominant Th1 response is a major cause of chronic non-healing disease where mice developing a Th2 response are more susceptible to *Leishmania* parasites (6).

During infection with *L. major* parasites, a strong Th1 immune response characterized by high levels of IFN-γ is associated with better parasite control (7–9). IFN-γ is necessary for control of *Leishmania* infection where mice lacking the IFN-γ receptor are unable to control *L. major* infection (7). Additionally, mice lacking IFN-γ are more susceptible to chronic *L. amazonensis* infection compared to control mice (9). Interestingly, during *L. major* infection, the requirement for IFN-γ signaling is mainly isolated to myeloid lineage cells (8). For instance, mice selectively lacking the IFN-γ receptor on macrophages results in uncontrolled lesion growth following infection with *L. major* (8). This is most likely because IFN-γ stimulates macrophages, the main cell type harboring *Leishmania* parasites, to produce effector molecules like nitric oxide (NO) and reactive oxygen species (ROS) which directly limit parasite growth (10–12). In addition, IFN-γ activates a number of other cell autonomous programs many of which have not been investigated during CL. For instance, IFN-γ stimulates the expression of guanylate binding proteins (GBPs) (13). GBPs are a part of the IFN-inducible dynamin family of GTPases that exert multifactorial roles during disease and infection (13). These proteins are characterized by their ability to bind and hydrolyze GTP, but GBPs also play other significant roles in cells such as disrupting pathogen membranes, inducing inflammasome activation, and promoting pathogen cell death (13, 14). Specifically, GBPs possess pattern recognition capabilities and exert antimicrobial effects through their GTPase activity against bacteria, virus, and parasites (15–19).

GBPs localize to both host cell membranes and target intracellular pathogen membranes during infections such as *Toxoplasma gondii* and *Shigella flexneri* infection, leading to membrane disruption, exposure of the pathogen to the intracellular environment, or pathogen death (15–21). Ultimately membrane disruption leads to the recruitment of supramolecular structures to activate cell death pathways, such as apoptosis or pyroptosis, through inflammasome formation and release of pro-inflammatory molecules (17). GBPs also aid in the recruitment of antimicrobial effectors to pathogen containing vacuoles, like NADPH oxidase subunits, and they facilitate the capture of intracellular microbes in autophagolysosomes (22–24). Additionally, GBPs can inhibit viral replication and suppress proteolytic activity of furin, a proprotein convertase which cleaves viral pro-proteins, further disrupting viral maturation (25, 26). Importantly, GBPs have been proposed as a new class of pattern recognition receptors binding directly to LPS, however additional ligands recognized by GBPs are still largely unknown (19, 27–29). GBPs also exert anti-microbial effects without localizing to the pathogen membrane (30, 31). For example, during *Chlamydia muridarum* infection, GBPs do not localize to the vacuole, but they are still required for rapid induction of pyroptosis in response to *Chlamydia* infection (30).

To date, minimal work has been done to understand the role of murine GBPs (mGBPs) during *Leishmania* infection. *mGbp2* and *mGbp5* are upregulated during infection with *L. major* in susceptible non-healing female Balb/c mice (32). Additionally, during *L. donovani* infection in non-phagocytic cells, IFN-γ upregulates GBP2 which is responsible for restriction of growth, despite GBP2 not localizing to the parasitophorous vacuole (PV) (31). Importantly, the expression of human GBP1 (hGBP1), hGBP2, hGBP4, and hGBP5 are elevated in human CL lesions (33, 34). However, the role of GBPs in cell autonomous defense against *Leishmania* species causing CL is incompletely understood leaving a wide knowledge gap. Here we demonstrate, the expression of GBPs is highly upregulated during CL. Moreover this work shows GBPs possess a transient role in controlling parasite burdens and disease severity in an experimental model of CL due to *L. major* infection. Ultimately, we find GBPs orchestrate the expression of inducible nitric oxide synthase (iNOS) in macrophages while selectively downregulating arginase (Arg-1), resulting in better control of *L. major* parasites. Unlike other infections, our data suggest that the function of GBPs during *L. major* infection is not dependent on localization to the pathogen, highlighting a unique feature of GBP biology in Leishmaniasis.

## Material and Methods

### mRNA extraction and real time PCR

mRNA was extracted with the Qiagen RNeasy Mini Kit 250 (Qiagen). RNA was reverse-transcribed with the High-Capacity cDNA reverse transcription kit (Applied Biosystems). Quantitative real-time PCR was performed using SYBR green PCR Master Mix and a QuantStudio 6 Flex real-time PCR system (Life Technologies). Mouse primer sequences were selected from the PrimerBank (http://pga.mgh.harvard.edu/primerbank/): Gbp2 (forward 5’- CTGCACTATGTGACGGAGCTA-3’ and reverse 5’- GAGTCCACACAAAGGTTGGAAA-3’), Gbp3 (forward 5’- GAGGCACCCATTTGTCTGGT-3’ and reverse 5’- CCGTCCTGCAAGACGATTCA-3’), Gbp5 (forward 5’- CAGACCTATTTGAACGCCAAAGA-3’ and reverse 5’- TGCCTTGATTCTATCAGCCTCT-3’), Gbp7 (forward 5’- TCCTGTGTGCCTAGTGGAAA-3’ and reverse 5’- CAAGCGGTTCATCAAGTAGGAT), and Gbp9 (forward 5’- GGTCACCGGGAATAGACTGG-3’ and reverse 5’- GGGCCACACTTGTCATAGCA-3’).

### Mice

Female C57BL/6NCr mice were purchased from the National Cancer Institute. GBP^Chr3^ KO mice were provided by Dr. Jörn Coers from Duke University and are previously described (35). Mice were used for experiments at 6 to 8 weeks of age and were housed under pathogen-free conditions at the University of Arkansas for Medical Sciences (UAMS). All procedures were approved by the UAMS IACUC and performed in accordance with institutional guidelines.

### Parasites

*Leishmania major* (WHO/MHOM/IL/80/Friedlin), *L. amazonensis* (MHOM/BR/00/LTB0016), or *L. mexicana* (MNYC/BZ/62/M379) parasites were grown *in vitro* in Schneider’s Drosophila medium (Gibco) supplemented with 20% heat-inactivated fetal bovine serum (FBS, Invitrogen), 2 mM L-glutamine (Sigma), 100 U/mL penicillin, and 100 μg/mL streptomycin (Sigma). Metacyclic promastigotes from 4-5-day old parasite cultures were isolated by Ficoll density gradient centrifugation (Sigma) (36).

### Generation of Bone Marrow-Derived Macrophages

Femurs collected from mice were soaked in 70% ethanol for 2 minutes and then flushed with 10 mL of cDMEM to extract bone marrow cells. Bone marrow cells were counted before plating 5×10^6^ cells per 100 mm Petri dish in 10 mL of conditioned macrophage media (cDMEM with 25% L929 cell supernatants). Cells were cultured for 7 days, refreshing media at day 3. To remove the macrophages from the Petri dish, macrophages were washed with ice-cold phosphate-buffered saline (PBS) and gently removed with a cell scraper. The collected macrophages were counted and loaded into 24-well plates with 1×10^6^ cells in 1 mL cDMEM per well.

### In vitro media supplementation

*In vitro* macrophage cultures were infected with *L. major* for 2 hours before cells were washed 2 times with PBS to wash away extracellular parasites. Cells were then cultured in media alone or media supplemented with 10 ng/mL IFN-γ (peprotech) or 100 ng/mL LPS (Sigma) + 10 ng/mL IFN-γ for 72 hours.

### Processing of cell cultures with cytospin analysis

At 72 hours post infection (hpi) samples were added to cytospin chambers and spun onto the slide at 1000 revolutions per minute (rpm) for 5 minutes using an automated cytospin (company). To quantify the number of parasites per macrophage, total uninfected and infected macrophages and total parasites were calculated manually on a brightfield upright microscope (Nikon). For each sample 100 infected macrophages were counted.

### Immunofluorescence microscopy

At 48 hpi, samples were fixed with 4% paraformaldehyde and 0.1% Triton-X before incubation with PBS and 0.5% BSA to block nonspecific binding. Samples were incubated with rabbit anti-GBP2 antibody (Proteintech) for one hour followed by goat anti-rabbit AF488 secondary antibody (Life Technologies) for 1 h. DAPI was added for 5 minutes and then coverslips with the samples were mounted onto slides with MOWIOL mounting media. Samples were visualized using a Ti2 eclipse microscope with a 60X oil immersion lens objective, and images were captured with a D5-QilMc digital camera (Nikon).

### Bulk RNA-Seq: Sample preparation

The Bulk RNA-Seq samples were prepared and data was acquired as a part of a previous study (37). Breifly, total RNA was isolated from the cell lysate of naive and infected ears by using Qiagen’s RNeasy plus mini kit according to the manufacturer’s instructions. The CTPR Genomics and Bioinformatics Core at the Arkansas Children’s Research Institute (ACRI) prepared sequencing libraries from RNA samples by use of the Illumina TruSeq Stranded mRNA Sample Preparation Kit v2. for sequencing on the NextSeq 500 platform using Illumina reagents. The quality and quantity of input RNA was determined using the Advanced Analytical Fragment Analyzer (AATI) and Qubit (Life Technologies) instruments, respectively. All samples with RQN (RNA quality number) values of 8.0 or above were processed for sequencing. Sequencing libraries were prepared by use of the TruSeq Stranded mRNA Sample Prep Kit (Illumina). Briefly, total RNA (500 ng) was subjected to polyA selection, then chemically fragmented and converted to single-stranded cDNA using random hexamer primed reverse transcription. The second strand was generated to create double-stranded cDNA, followed by fragment end repair and addition of a single A base on each end of the cDNA. Adapters were ligated to each fragment end to enable attachment to the sequencing flow cell. The adapters also contain unique index sequences that allow the libraries from different samples to be pooled and individually identified during downstream analysis. Library DNA was PCR amplified and enriched for fragments containing adapters at each end to create the final cDNA sequencing library. Libraries were validated on the Fragment Analyzer for fragment size and quantified by use of a Qubit fluorometer. Equal amounts of each library were pooled for sequencing on the NextSeq 500 platform using a high output flow cell to generate approximately 25 million 75 base reads per sample.

### RNASeq Analysis

Data analysis was performed for a previous study (37). Following demultiplexing, RNA reads were checked for sequencing quality using FastQC (http://www.bioinformatics.babraham.ac.uk/projects/fastqc) and MultiQC (38)(version 1.6). The raw reads were then processed according to Lexogen’s QuantSeq data analysis pipeline with slight modification. Briefly, residual 3’ adapters, polyA read through sequences, and low quality (Q < 20) bases were trimmed using BBTools BBDuk (version 38.52) (https://sourceforge.net/projects/bbmap/). The first 12 bases were also removed per the manufacture’s recommendation. The cleaned reads (> 20bp) were then mapped to the mouse reference genome (GRCm38/mm10/ensemble release-84.38/ <u>GCA_000001635.6</u>) using STAR (39) (version 2.6.1a), allowing up to 2 mismatches depending on the alignment length (e.g. 20-29bp, 0 mismatches; 30-50bp, 1 mismatch; 50–60+bp, 2 mismatches). Reads mapping to > 20 locations were discarded. Gene level counts were quantified using HTSeq (htseq-counts) (40) (version 0.9.1) (mode: intersection-nonempty).

Genes with unique Entrez IDs and a minimum of ∼2 counts-per-million (CPM) in 4 or more samples were selected for statistical testing. This was followed by scaling normalization using the trimmed mean of M-values (TMM) method (41) to correct for compositional differences between sample libraries. Differential expression between naive and infected ears was evaluated using limma voomWithQualityWeights (42) with empirical bayes smoothing. Genes with Benjamini & Hochberg (43) adjusted p-values ≤ 0.05 and absolute fold-changes ≥ 1.5 were considered significant.

Gene Set Enrichment Analysis (GSEA) was carried out using Kyoto Encyclopedia of Genes and Genomes (KEGG) pathway databases and for each KEGG pathway, a p-value was calculated using hypergeometric test. Cut-off of both p < 0.05 and adjusted p-value/FDR value < 0.05 was applied to identify enriched KEGG pathways. DEGs that are more than 1.5-fold in *L. major*-infected ears relative to uninfected controls were used as input, with upregulated and downregulated genes considered separately. Subsequently, the heat maps were generated using these genes with complex Heatmap. All analyses and visualizations were carried out using the statistical computing environment R version 3.6.3, RStudio version 1.2.5042, and Bioconductor version 3.11. The raw data from our bulk RNA-Seq analysis were deposited in Gene Expression Omnibus (GEO accession number—GSE185253).

### In vivo infections

For infections, 100,000 (Figure 1) or 2 x 10^6^ (all other figures) parasites were intradermally injected into the right ear in a volume of 10 μL of PBS (Gibco). The contralateral left ear was not injected with parasites, serving as an uninflamed control. Ear thickness and lesion diameters were recorded weekly with electronic calipers and lesion volume was calculated. Ears were digested enzymatically for 90 minutes at 37°C in 0.25 mg/mL liberase (Roche) with 10 μg/mL DNase I (Sigma) in RPMI 1640 (Gibco). Tissue parasite burden was determined using limiting dilution assays (44).

**Figure 1:**
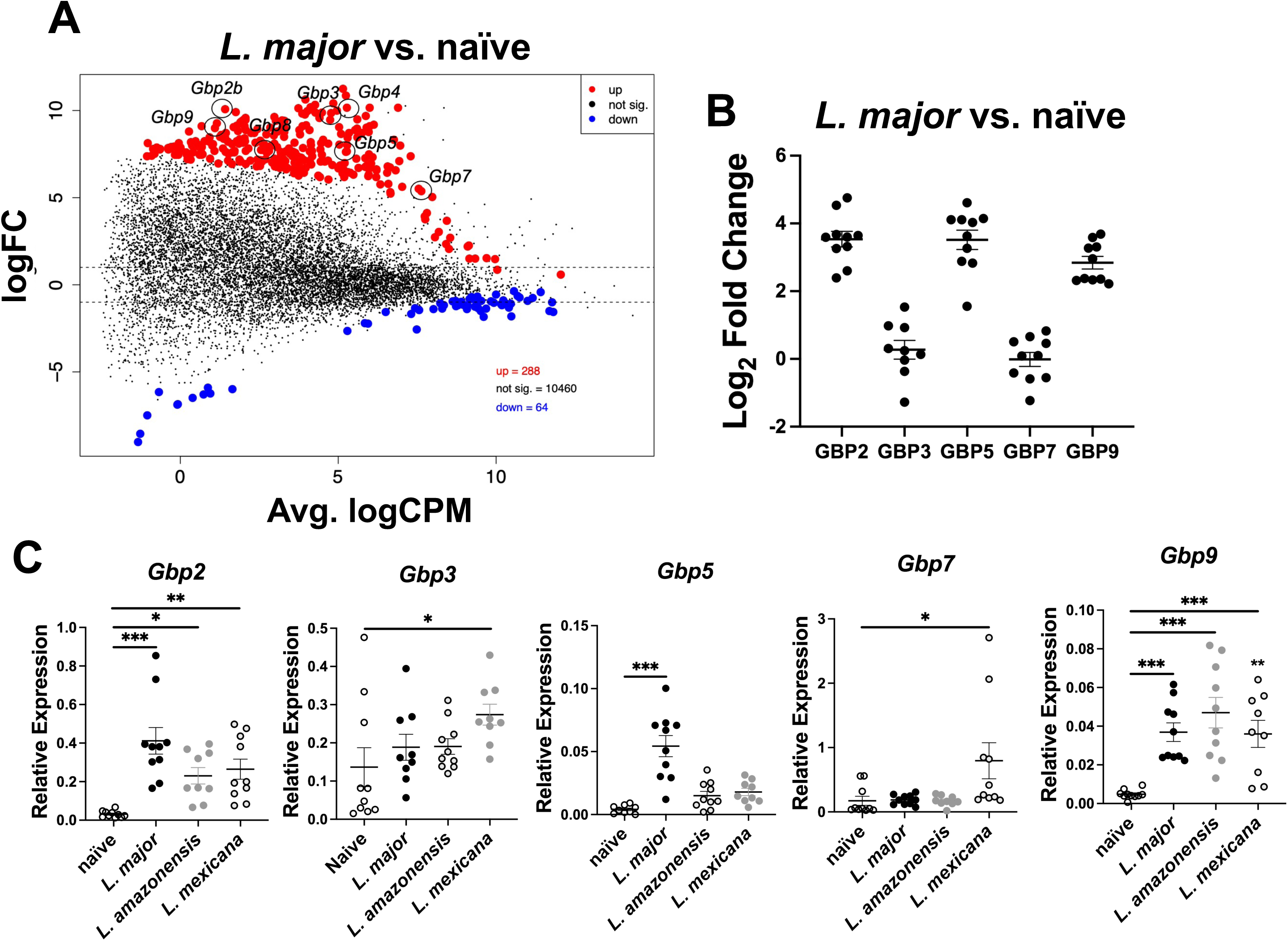
GBP expression is a hallmark of CL. Mice were infected intradermally with 2 x 10^6^ *L. major* parasites and at 4 weeks post infection (wpi) RNA was isolated from ear tissue and prepped for bulk RNA-Seq. (A) An MDS plot depicting up and down regulated transcripts during *L. major* infection. Red transcripts are upregulated with infection and blue transcripts are downregulated with infection. (B) Real-time PCR (RT-PCR) was conducted on RNA isolated from ear lesions from naïve mice or mice infected with 100,000 *L. major* parasites. Expression of GBPs during *L. major* infection was compared against the expression of each respective GBP in naïve mice to identify the fold change. (C) RT-PCR was performed on RNA isolated from ear lesions from mice infected with either 100,000 *L. major*, *L. amazonensis*, or *L. mexicana* at 6 wpi. Data in (B-C) is pooled from two independent experiments with 5 mice per group and results are shown as mean ± SEM. Significance was determined using a t-test compared to the naïve samples where *p<0.05 **p<0.01, ***p<0.001. Significance is shown in comparison to naïve samples.

### Single-cell RNASequencing Sample Preparation

The scRNASeq samples were prepared and data was acquired as a part of a previous study (37). In short, the Arkansas Children’s Research Institute (ACRI) Genomics and Bioinformatics Core prepared NGS libraries from fresh single-cell suspensions using the 10X Genomics NextGEM 3’ assay for sequencing on the NextSeq 500 platform using Illumina SBS reagents. Trypan Blue exclusion determined cell quantity and viability. Library quality was evaluated with the Advanced Analytical Fragment Analyzer (Agilent) and Qubit (Life Technologies) instruments.

### scRNASeq Data Analysis

Data analysis was performed as a part of a previous study (37). Briefly, the UAMS Genomics Core generated Demultiplexed fastq files which were analyzed using 10X Genomics Cell Ranger alignment and gene counting software, a self-contained scRNASeq pipeline developed by 10X Genomics. The reads were aligned to the mm10 reference transcriptomes using STAR and transcript counts were generated (39, 45). The *Seurat* R package processed the raw counts generated by *cellranger count* (46, 47). Potential doublets, low quality cells, and cells with a high percentage of mitochondrial genes were filtered out. Cells that have unique feature counts > 75^th^ percentile plus 1.5 times the interquartile range (IQR) or < 25^th^ percentile minus 1.5 time the IQR were filtered. Similarly, cells with mitochondrial counts falling outside the same range for mitochondrial gene percentage were filtered. After filtering, all 8 sequencing runs were merged. The counts were normalized using the LogNormalize method which log-transforms the results (37). Subsequently, the 2000 highest variable features were selected. The data was scaled, and Principal component analysis (PCA) was performed. A JackStraw procedure was implemented to determine the significant PCA components that have a strong enrichment of low p-value features.

A graph-based clustering strategy embedded cells in graph structure (48). Seurat visualized the results in t-distributed stochastic neighbor embedding (tSNE) and Uniform Manifold Approximation and Projection (UMAP) plots (49). Seurat *FindNeighbors* and *FindClusters* functions were optimized to label clusters. Seurat *FindAllMarkers* function finds markers that identify clusters by differential expression, defining positive markers of a single cluster compared to all other cells and comparing those to known markers of expected cell types from previous single-cell transcriptome studies. Cell type identifications were determined by manually reviewing these results, and some clusters were combined if their expression was found to be similar. From here for this work, we specifically provide Feature maps showing transcript expression of GBP2, GBP3, GBP5, GBP7, and GBP9 amongst all clusters, and particularly in macrophages and monocytes.

### Flow cytometry

Flow cytometric analysis was conducted *in vivo* on ear tissue. For *in vivo* flow cytometric analysis, ear tissue was enzymatically digested and processed for flow cytometric analysis of dermal cells from the ears. To determine cellular viability, cells were incubated with a Zombie Aqua viability dye (Biolegend). Fc receptors were blocked with 2.4G2 anti-mouse CD16/CD32 (Invitrogen or BioXCell) and 0.2% rat IgG (BioXCell). Cells were surfaced stained with antibodies against CD45 BV650 (BD Horizon) or AF700 (eBioscience), CD11b BV750 (Biolegend) or BV605 (Biolegend), CD64 BV421 (Biolegend) or BV711 (Biolegend), Ly6C PercpCy5.5 (Invitrogen), Ly6G AF700 (Biolegend). Intracellular stain was performed using the Foxp3 Transcription Factor Staining Buffer kit (Invitrogen). Intracellular molecules and cytokines were stained with antibodies against Arg-1 PE (Invitrogen) or iNOS APC (Biolegend). Cell events were acquired on a Cytek Northern Lights (Cytek) and analyzed using FlowJo (Tree star).

### Statistics

All data were analyzed for statistical significance using GraphPad Prism 9. Statistical significance was determined using a two-tailed Student’s unpaired t-test or a two-way anova with a Tukey’s multiple comparison test. Outliers were identified by a Grubb’s outlier test and removed.

## Results

GBPs orchestrate the host defense to many intracellular pathogens. To determine if GBPs are involved in the host defense against *Leishmania* parasites we analyzed the expression of GBPs during *in vivo L. major* infection. Briefly, C57BL/6 mice were intradermally infected with *L. major* parasites in the ear and at 4 weeks post-infection (wpi) RNA-Sequencing was conducted on whole lesions from mice infected with *L. major* and compared to naïve controls (Figure 1A). Multiple GBPs were upregulated with *L. major* infection including *Gbp2*, *Gbp3*, *Gbp4*, *Gbp5*, *Gbp7*, *Gbp8*, and *Gbp9* (Figure 1A). To validate our RNA-seq findings and identify the most highly upregulated GBPs during *L. major* infection, we performed real-time PCR (RT-PCR) to quantify GBP expression relative to naïve samples. By utilizing Log₂ fold change as a comparative metric we can prioritize candidates for further investigation. GBP2, GBP5, and GBP9 were most upregulated with *L. major* infection compared to naïve samples (Figure 1B). After confirming that GBP expression is associated with *L. major* infection, we next determined whether this upregulation is specific to *L. major* or also occurs in response to infections with other *Leishmania* species. Mice were infected with either *L. major*, *L. mexicana*, or *L. amazonensis* parasites intradermally in the ear. Following establishment of infection and lesion formation, ear tissue was collected and prepped for RT-PCR (Figure 1C). Interestingly, *Gbp2* and *Gbp9* were also elevated following infection with both *L. mexicana* and *L. amazonensis*, while *Gbp5* trended towards an increase following infection with both species similar to *L. major* infection (Figure 1C). The expression of *Gbp3* and *Gbp7* increased following infection with *L. mexicana* (Figure 1C). These data suggest that multiple GBPs are elevated *in vivo* after infection with different *Leishmania* species, and that GBP expression is a hallmark of CL, although different GBPs may be elevated depending on the species of infecting parasite.

Next, to investigate whether GBPs play a role in host defense during *L. major* infection, we set up an *in vitro* assay to assess parasite burdens in BMDMs as macrophages are the main cell type harboring and killing parasites during *Leishmania* infection. Specifically, macrophages were derived from control C57BL/6 mice or GBP^Chr3^ KO mice which are deficient for all the GBPs contained on Chromosome 3 including GBP1, GBP2, GBP3, GBP5, and GBP7 that has been described previously (35). GBPs often work in concert with one another and the deletion of only one or two GBPs may lead to compensation by other GBPs. Deletion of GBPs contained on chromosome 3 has shown efficacy in studying GBPs role in health and disease (50–53). These GBPs are more often associated with roles in the immune response (20, 24, 25, 50). To analyze parasite burdens in macrophages with and without GBPs we differentiated macrophages from bone marrow cells from either control or GBP^Chr3^ KO mice. After differentiation, BMDMs were infected with *L. major*, or infected and then treated with IFN-γ, or IFN-γ and LPS. Cytospin was used to calculate parasite burdens (Figure 2A-B). We observed no significant differences between parasites per macrophage or the percent infectivity of control or GBP^Chr3^ KO macrophages during infection with *L. major* after 72 hours of culture in media (Figure 2B). However, upon treatment with IFN-γ, GBP^Chr3^ KO macrophages harbored significantly more parasites per macrophage (Figure 2A-B). Additionally, the percent of infected macrophage was higher for GBP^Chr3^ KO macrophages compared to controls that were infected and treated with IFN-γ (Figure 2A-B). There were no differences in parasite burden in control or GBP^Chr3^ KO macrophages infected and treated with LPS/IFN-γ suggesting the defect in parasite control can be overcome depending on the external stimuli (Figure 2B). These data suggest that upon stimulation with IFN-γ, GBPs play a role in parasite control, and during GBP deficiency macrophages are less equipped to combat the infection.

**Figure 2:**
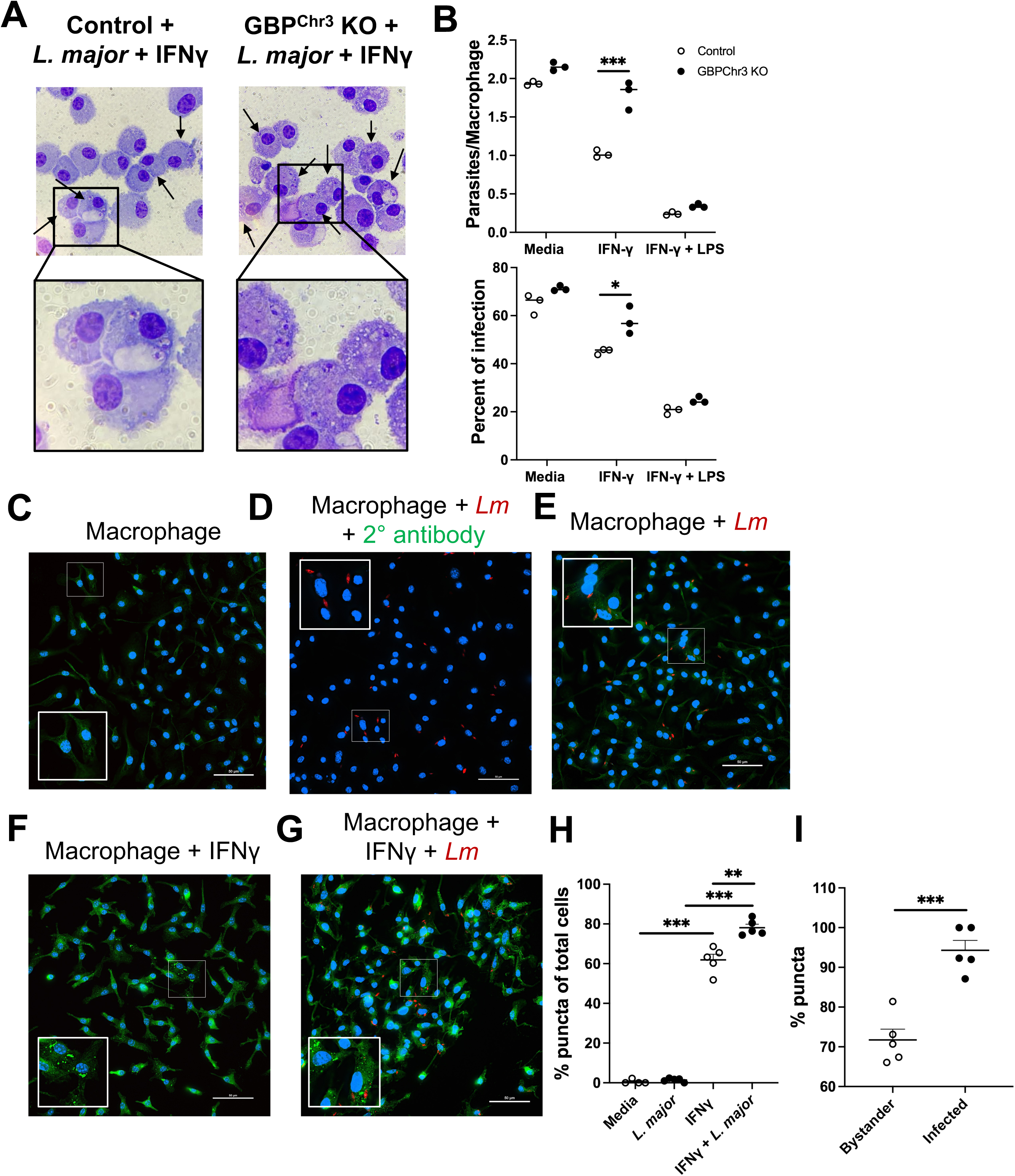
Macrophage GBPs control *L. major* parasites independent of localization to the parasite membrane. BMDMs from control mice or GBP^Chr3^ KO mice were infected and cultured in media or different supplemented media conditions before cells were prepped for cytospin analysis or immunofluorescence microscopy. (A) Representative images of control or GBP deficient macrophages infected with *L. major* and treated with IFN-γ for 72 hours. (B) Quantification of (A) showing parasites per macrophages and percent of infection after infection alone, infection and treatment with IFN-γ, or infection and treatment with LPS + IFN-γ after 72 hpi. (C-G) After 48 hpi, cells were fixed and stained with an antibody against GBP2 for immunofluorescence microscopy. Representative images of macrophages (C) cultured in media, (D) infected with *L. major* and only stained with a secondary antibody, (E) infected with *L. major* and stained with an antibody against GBP2, (F) IFN-γ-treated macrophages, or (G) IFN-γ-treated macrophages infected with *L. major*. (H) Quantification of the percentage of puncta containing cells as a percentage of total cells in (C-G) and (I) quantification of bystander versus infected cells containing puncta in (G). Data are representative of 2 or 3 independent experiments with at least 3 mice per group. Significance was determined using a two-way ANOVA paired with a Tukey’s multiple comparison test where *p<0.05 **p<0.01, ***p<0.001.

GBP localization to vacuole membranes is intimately tied to function and pathogen control (54, 55). Therefore, we investigated whether GBPs localizes to *L. major* to determine if GBP function during *L. major* infection is tied to cellular localization. We chose to analyze GBP2 protein based on our transcriptomic data where GBP2 is the most highly elevated during infection with *L. major* compared to the other GBPs (Figure 1). To analyze GBP2 localization during *L. major* infection, BMDMs were cultured in media and either left uninfected, infected with *L. major*, treated with IFN-γ, or infected and treated with IFN-γ. At 48 hpi, samples were stained with an antibody against GBP2. First, to determine the expression of GBP2 during homeostasis, GBP2 expression was analyzed in macrophages cultured in media alone (Figure 2C). We detected a diffuse signal of GBP2 within uninfected untreated cells (Figure 2C). Importantly, the signal detected in untreated macrophages was not due to background of the secondary antibody. When untreated macrophages were stained with the secondary antibody alone, there was no diffuse signal detected within the cells suggesting that during homeostasis GBP2 is expressed ubiquitously throughout the cell (Figure 2D). During infection of untreated macrophages, we observed similar GBP2 expression compared to uninfected BMDMs cultured in media alone (Figure 2E). However, upon treatment with IFN-γ, GBP2 was highly expressed and detected throughout the cell, manifesting mostly as puncta (Figure 2F). The GBP2 localization during treatment with IFN-γ in the absence of parasites was spread out throughout the cell (Figure 2F). Interestingly, upon *L. major* infection and IFN-γ treatment, GBP2 was similarly localized throughout the cell compared to treatment with IFN-γ alone (Figure 2G). Infected and uninfected bystander macrophages stimulated with IFN-γ possessed punctate GBP2 throughout the cell (Figure 2G). Importantly, we did not detect GBP2 signal localized to the parasite membrane although many GBP puncta were detected in close association with the parasite signal. These data suggest macrophage activation leads to increased GBP2 protein, however although GBPs contribute to control of *L. major* parasites, this activity is not achieved through GBP2 localization to the intracellular parasite.

We next measured the percentage of cells with GBP2 puncta within each condition. We only detected GBP puncta in infected, bystander, and uninfected cells that were treated with IFN-γ and not in macrophages with no pro-inflammatory stimulus (Figure 2G-H). Interestingly, we found that BMDMs treated with both IFN-γ and *L. major* possessed more puncta-positive cells than those treated with IFN-γ alone (Figure 2H). Next, we investigated which cells possess puncta, specifically infected cells or in nearby uninfected bystander cells during treatment with IFN-γ and infection with *L. major*. Interestingly, many bystander cells contained GBP2 puncta (Figure 2I). However, the percentage of puncta-positive cells was significantly higher among infected cells compared to bystander uninfected cells (Figure 2I). These data together suggest expression of GBP2 puncta is somewhat specific to cells that contain *L. major*.

Because GBP expression is increased with infection, and IFN-γ treated GBP^Chr3^ KO macrophages harbor more parasites compared to control macrophages, we next investigated if GBPs play a role in parasite control or disease pathology during *in vivo L. major* infection. To analyze GBPs during *in vivo L. major* infection, we utilized the previously mentioned mouse strain deficient for the GBPs located on chromosome 3. To initially determine if GBPs are important to host defense during *L. major* infection, control or GBP^Chr3^ KO mice were intradermally infected with *L. major* parasites and lesion development was monitored weekly (Figure 3A). Interestingly, GBP^Chr3^ KO mice exhibit significantly larger lesions at 2 wpi compared to control infected mice where both the thickness and volume is significantly larger than controls (Figure 3A-B). Furthermore, we analyzed local and systemic parasite burdens and found that GBP^Chr3^ KO mice possess higher local parasite burdens in the skin compared to control mice (Figure 3C). Systemic parasite burdens in the draining lymph node (dLN) and spleen were unchanged compared to control mice (Figure 3C and data not shown). These data suggest GBPs play a role in disease control during *L. major* infection and in the absence of GBPs, parasite burden is elevated, resulting in worse disease severity.

**Figure 3:**
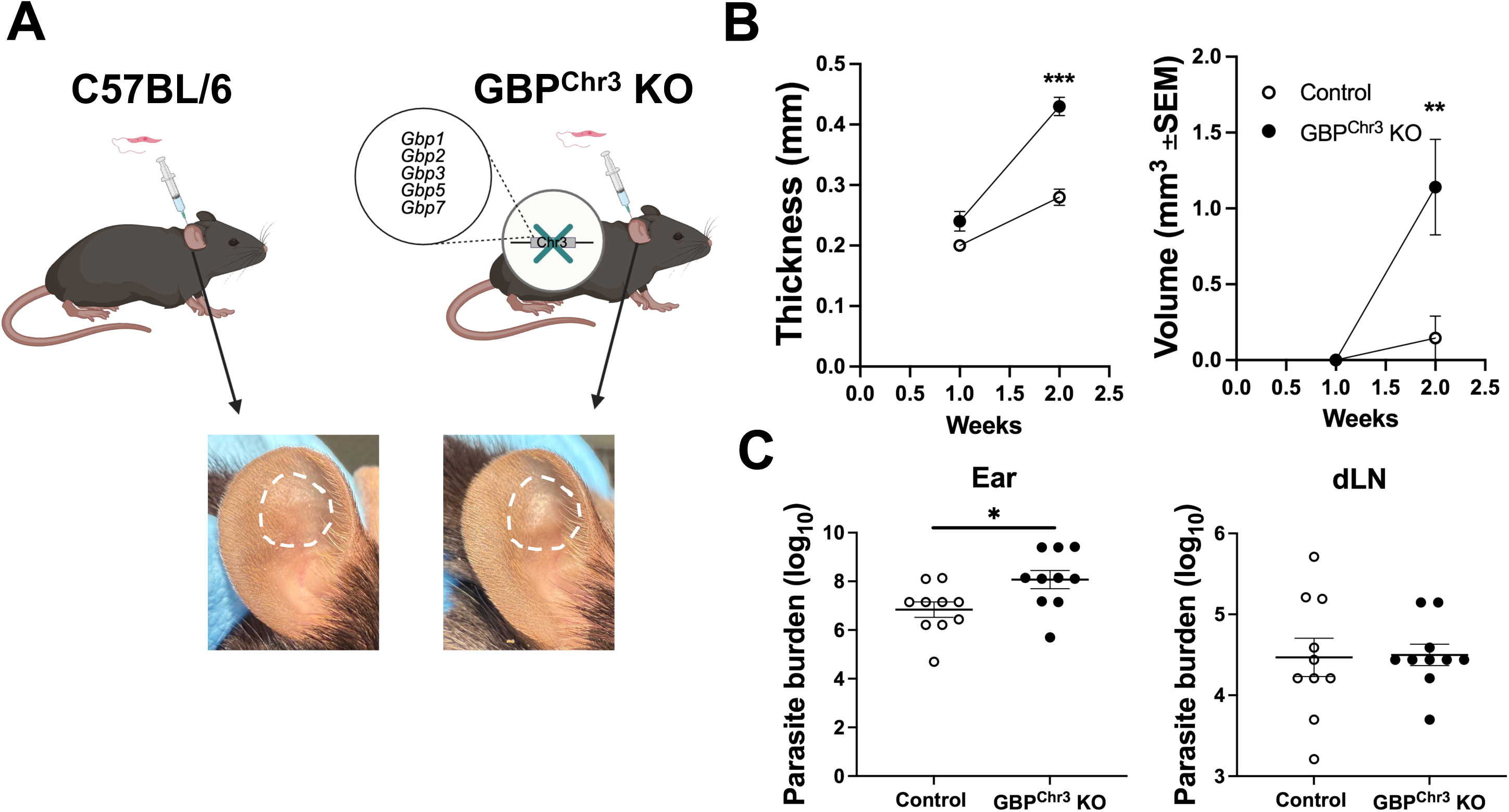
GBPs participate in control of parasites and disease severity during *L. major* infection. C57BL/6 control or GBP^Chr3^ KO mice were infected intradermally with 2 x 10^6^ *L. major* parasites. (A) Diagram showing infection of control or GBP^Chr3^ KO mice and representative lesions from each mouse strain at 2 wpi. (B) Lesions were monitored weekly by measuring ear thickness and lesion volume with electronic calipers. (C) The local ear and systemic dLN parasite burdens were quantified using a limiting dilution assay (LDA). Data is pooled from 3 experiments with 5 mice per group per experiment. Significance was determined using a t-test where *p<0.05 **p<0.01, and ***p<0.001.

Because GBP expression is necessary for optimal parasite control, we next analyzed the cell types expressing GBPs during *in vivo L. major* infection to investigate the cells being affected by GBP deletion. We hypothesized GBPs are expressed in myeloid derived cells during *L. major* infection as these are the cells responsible for harboring and killing parasites. Additionally, our previous data determined BMDMs deficient in GBPs on chromosome 3 harbor significantly more parasites compared to control BMDMs (Figure 2A-B). To determine which cell types express GBPs during infection, mice were infected with *L. major* parasites and at 4 wpi tissue was taken for Single-cell RNA Sequencing (scRNA-seq) from naïve skin and CL lesions as a part of a previous study (37). The scRNA-seq data revealed 35 distinct cell populations in the lesion present after *L. major* infection (Figure 4A). We analyzed the expression of *Gbp2*, *Gbp3*, *Gbp5*, *Gbp7*, and *Gbp9* and the findings are consistent between each GBP. In the naïve uninfected skin tissue, resident macrophages are the main cell type expressing GBPs (Figure 4B-C). Interestingly, during *L. major* infection, macrophages, particularly monocyte-derived macrophages and resident macrophages, are the main cell types expressing GBPs (Figure 4B-C). Monocyte-derived macrophages originate from inflammatory monocytes recruited to the infection site to replenish resident macrophages. We find iMonos also exhibit elevated GBP expression with *L. major* infection, but to a lesser extent than monocyte-derived macrophages and resident macrophages. This is relevant as both resident macrophages, monocyte-derived macrophages, and iMonos become infected with *L. major* and are responsible for either serving as a permissive niche facilitating infection or as a resistant host cell orchestrating parasite killing. Additionally, fibroblast, T cells, and neutrophils express GBPs during *L. major* infection. Altogether, these data highlight the preferential, yet unrestricted expression of GBPs in myeloid cells.

**Figure 4:**
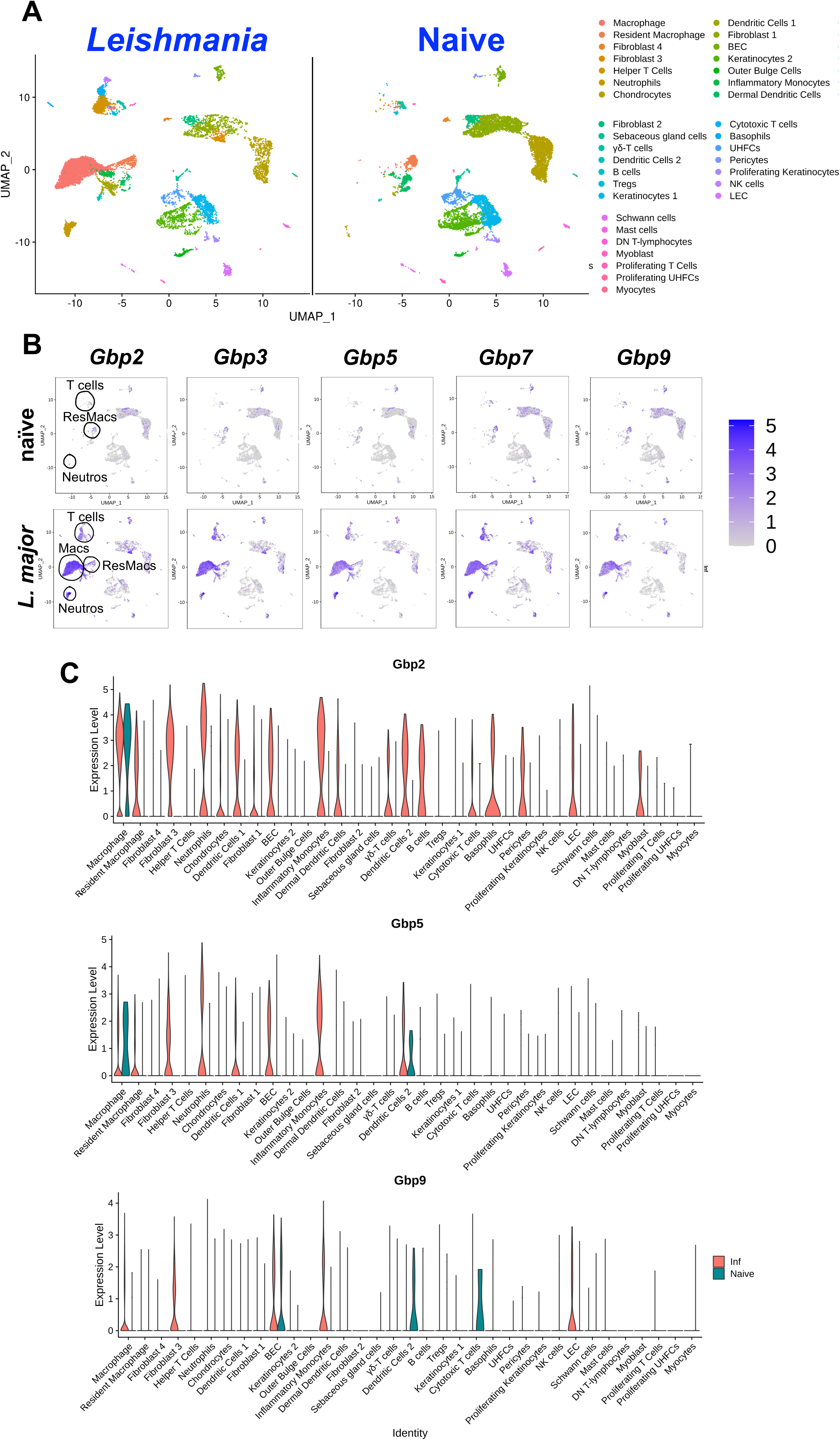
GBP expression is associated with dermal resident macrophage and monocyte-derived macrophage populations during *L. major* infection. Mice were infected with *L. major* and at 4 wpi, scRNA-seq was performed on naïve ear skin and CL lesions. (A) UMAP plots depict 35 unique cell populations were identified in the ear dermis during *L. major* infection. (B) Feature plots of expression distribution for *Gbp2*, *Gbp3*, *Gbp5*, *Gbp7* and Gbp9 in naïve animals and *L. major* infected mice. Expression levels for each gene are color-coded and overlaid onto UMAP plot. Cells with the highest expression level are colored dark purple. (C) Differential expression of selected GBP transcripts in 35 different cell types.

After determining macrophages are the main cell type expressing GBPs during *L. major* infection, we investigated macrophages from WT and GBP^Chr3^ KO mice during *L. major* infection. Given macrophages most highly express GBPs during *L. major* infection, we hypothesized the macrophage population is responsible for the difference in parasite burden in GBP^Chr3^ KO mice compared to controls. Additionally, because we did not observe GBP localization to *L. major* in vitro, we hypothesize the difference in parasite burden between control and GBP^Chr3^ KO mice was attributed to an alternative mechanism of control by GBPs. To further understand the role of GBPs in parasite control we analyzed the cellular phenotype of macrophages during *L. major* infection in mice with and without GBPs. Specifically, we employed flow cytometry to assess macrophage inducible nitric oxide (iNOS) and arginase-1 (Arg-1) levels. We chose to analyze these two molecules because iNOS expression is generally associated with NO production and parasite control, while Arg-1 is associated with susceptible hosts and parasite survival in leishmaniasis. Specifically, Arg-1 leads to the synthesis of polyamines which directly contributes to parasite replication (56). So, iNOS and Arg-1 positivity is used as a surrogate to probe between M1 versus M2 macrophages within tissues. To assess macrophage phenotype, mice were infected intradermally with *L. major* parasites and at 2 wpi, ear tissue was subjected to flow cytometry.

We assessed the percentage and total number of iNOS^+^ macrophages and iMonos. We identified a significant decrease in the percentage of iNOS^+^ macrophages in lesions in GBP^Chr3^ KO mice compared to control mice (Figure 5A-B). We observed a similar effect for iMonos as well where GBP^Chr3^ KO mice exhibited a lower percentage of iNOS^+^ iMonos compared to controls (Figure 5C-D). Additionally, not only was the iNOS^+^ populations of macrophages and iMonos reduced, but we also observed a significant increase in the percentage of macrophages and iMonos that displayed Arg-1 positivity (Figure 5E-H). Furthermore, the expression ratio of iNOS^+^ cells to Arg-1^+^ cells was significantly decreased for both macrophages and iMonos (Figure 5I-J). These data suggest myeloid lineage cells within GBP^Chr3^ KO are more characteristic of M2-like cells which are associated with parasite survival and disease progression. Together these data suggest that immune cells from GBP^Chr3^ KO mice possess an enhanced Arg-1 M2 program. Importantly, during GBP deficiency and *L. major* infection, the lesional macrophages and iMonos display a parasite friendly environment, consistent with higher parasite burdens in the GBP^Chr3^ KO mice compared to control mice.

**Figure 5:**
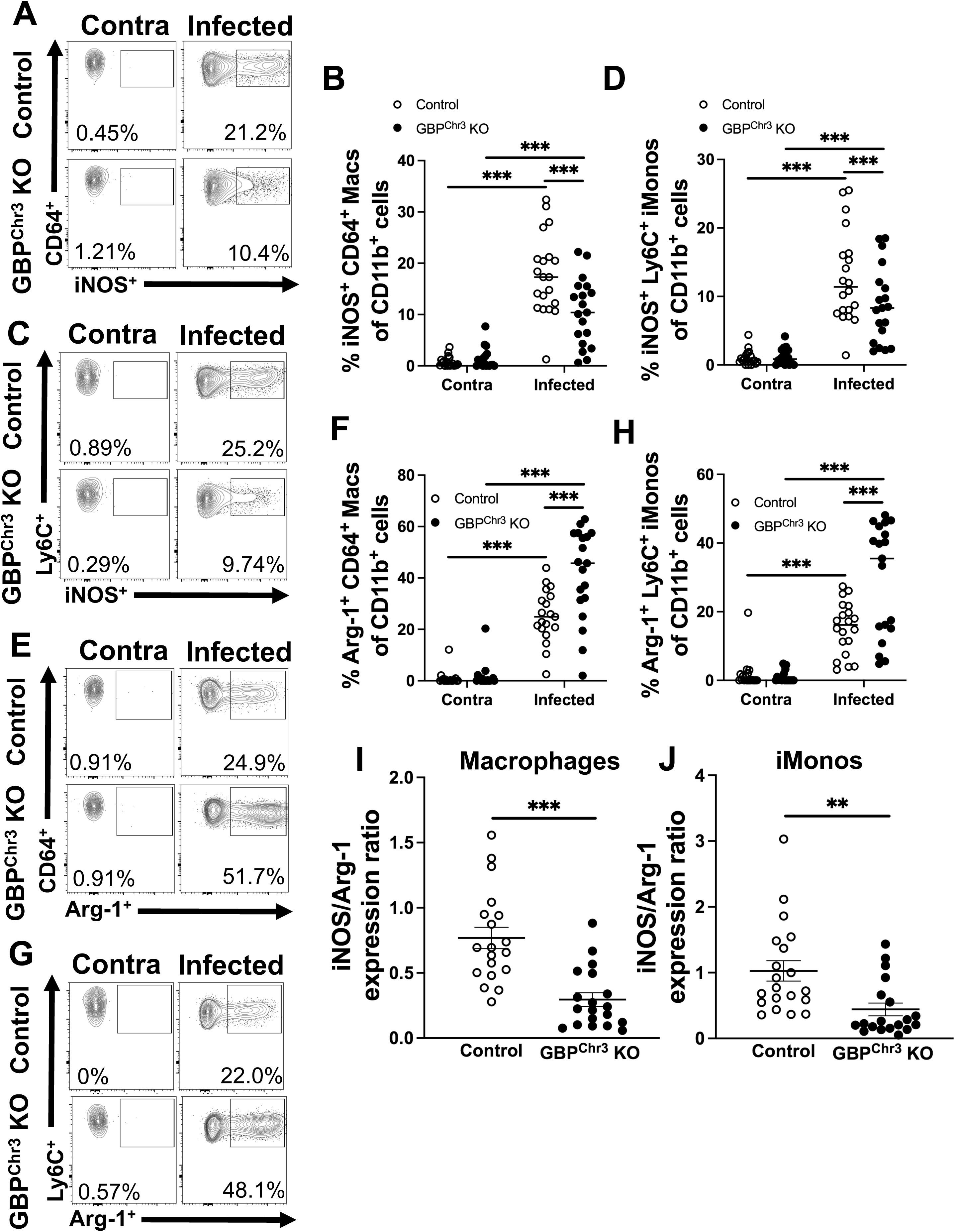
GBP deficiency results in reduced iNOS^+^ myeloid cells and increased Arg-1^+^ myeloid cells. Mice were infected with *L. major* parasites and at 2 wpi flow cytometric analysis was performed on ear tissue from control or GBP^Chr3^ KO mice. (A) Representative flow plots of iNOS^+^ macrophages are shown. Gated on live, single, CD45^+^, CD11b^+^, CD64^+^, Ly6G^-^ cells. (B) Quantification of (A) shows percentage and number of iNOS^+^ macrophages in the infected and contralateral ear of control or GBP^Chr3^ KO mice. (C) Representative flow plots of iNOS^+^ iMonos are shown. Gated on live, single, CD45^+^, CD11b^+^, Ly6C^+^, Ly6G^-^ cells. (D) Quantification of (C) shows percentage and number of iNOS^+^ iMonos in the infected and contralateral ear of control or GBP^Chr3^ KO mice. (E) Representative flow plots of Arg-1^+^ macrophages are shown. (F) Quantification of (E) showing percentage and number of Arg-1^+^ macs. (G) Representative flow plots of Arg-1^+^ iMonos are shown. (H) Quantification of (G) showing percentage and number of Arg-1^+^ iMonos. (I) expression ratio of iNOS^+^ macrophages to Arg-1^+^ macrophages. (J) Expression ratio of iNOS^+^ iMonos to Arg-1^+^ iMonos. Data are pooled from 3 experiments with 5-10 mice per group per experiment. Data are shown as mean. Significance was determined using a two-way ANOVA paired with a Tukey’s multiple comparison test where *p<0.05 **p<0.01, ***p<0.001.

## Discussion

GBPs are involved in the host response to a wide variety of intracellular pathogens (19). In this study, we demonstrate for the first time that GBPs play a crucial role in host defense and parasite control during an *in vivo* experimental model of *Leishmania* infection. Specifically, we found that the expression of GBPs is a hallmark of CL. Various GBPs were upregulated following infection with *L. major*, *L. mexicana*, and *L. amazonensis*. Previous work found that infection with *L. major* in mice of different genetic backgrounds is associated with the expression of Gbp2 and Gbp5 (32). However, this prior work did not establish a direct role for GBPs in parasite control, as we have shown here. Moreover, it was previously determined that during *L. major* infection, GBP2 is highly expressed in the skin tissue of semi-resistant mouse models within infected cells (32). In contrast, in the highly susceptible Balb/c mouse strain, GBP expression is decreased, and less infected cells express GBPs (32). This suggests that GBPs may influence disease outcome, with more favorable disease outcomes occurring when GBPs are highly expressed in cells infected with *Leishmania* parasites. Whether this holds true for infections with *Leishmania* species that cause more severe disease outcomes, such as *L. amazonensis* and *L. mexicana*, remains to be determined and is an active area of investigation in the lab.

In this work we define the loss of GBPs during infection with *L. major* parasites increases parasite burdens compared to controls *in vivo*. We speculate this is due to an increase in the percentage of Arg-1^+^ CD64^+^ macrophages and a decrease in iNOS^+^ CD64^+^ macrophages, reflecting a phenotypic shift in the macrophage population. The iNOS/Arg-1 ratio is a key indicator of whether the environment is permissive or resistant to parasites (57, 58). Specifically, cases of *Leishmania* infection characterized by a robust M2 macrophage response are associated with worse disease outcomes and uncontrolled parasite replication (57, 59, 60). For example, during *L. major* infection in both C57BL/6 and Balb/c mice, an Arg-1^+^ population is present (61). However, in susceptible Balb/c mice arginase expression begins around lesion formation and progressively increases until the experimental endpoint (61). In contrast, during infection in resistant C57BL/6 mice, arginase expression peaks at 3 wpi, when lesions are present, and then gradually declines as the lesion resolves (61).

We found that macrophages from GBP^Chr3^ KO mice exhibit lower percentages of iNOS positivity, suggesting that in the absence of GBPs, a more parasite-permissive environment is created. We suspect this is likely due to impaired iNOS-mediated parasite killing. Supporting this, during infection with the Seidman non-healing strain of *L. major*, a population of M2 mannose receptor-high dermal macrophages propagates infection, leading to non-healing lesions (60). Notably, iNOS positivity is not limited to infected cells (62). In fact, recruited monocytes rapidly induce iNOS, with the percentage of iNOS^+^ cells exceeding the percentage of infected cells (62). Importantly, the rapid induction of iNOS was linked to both TNF-α and IFN-γ production (62). Whether these factors are intact in the skin during infection in GBP-deficient mice remains to be determined. Furthermore, we detected elevated levels of activated TNF-α-positive CD4^+^ and CD8^+^ T cells in the dLNs, suggesting that the T cell compartment may be compensating for the reduction in iNOS expression in myeloid cells (data not shown). Taken together, these data suggest that during infection with *L. major*, GBPs orchestrate the NO gradient, which is crucial for subsequent parasite control.

Interestingly, by utilizing a photoconvertible pathogen labeling system, Müller et al., determined iNOS directly influences parasite growth by suppressing *Leishmania* metabolism (63). Mice deficient in iNOS and infected with *L. major* display increased lesion sizes compared to control mice starting at 4 wpi and continuing through 10 wpi (64). This suggests that, although GBP^Chr3^ KO mice exhibit a defect in the iNOS^+^ cellular compartment, their phenotype does not entirely match that of iNOS-deficient mice. We observed significant differences in lesion sizes between GBP^Chr3^ KO mice and controls at 2 and 3 wpi. However, the difference in lesion size between groups disappeared by 4 wpi, with GBP^Chr3^ KO mice ultimately healing their lesions in a manner similar to control mice (data not shown). This indicates that, despite an initial defect in parasite control during GBP^Chr3^ deficiency, the host can overcome differences in parasite burdens, suggesting compensation by another mechanism after 4 wpi. However, whether GBP^Chr3^ deficiency affects disease progression in more severe cases of CL, such as infection with *L. amazonensis* or *L. mexicana*, remains unknown and is an ongoing area of research in our lab.

Previous work showed that during *T. gondii* infection, which stimulates GBP recruitment to the PV, GBP2^+^ PVs are highly enriched for iNOS activity (54). In this context, GBP2 and iNOS appear to work in concert, with inhibition of iNOS leading to the shedding of GBP2 from the PV and increased parasite replication (54). Ultimately, iNOS was found to be necessary for control of *T. gondii*, suggesting that GBPs on Chr3 play a key role in orchestrating optimal iNOS-mediated parasite clearance (54). In our study, we determined that GBPs do not localize to *L. major* during *in vitro* infection (Figure 2C-G). Specifically, we found that following IFN-γ treatment, GBPs are expressed throughout the cell and upon infection with *L. major*, GBP localization does not change and GBP2 does not migrate to *L. major*. Consistent with our findings, previous research with *L. donovani* infection found that GBPs do not localize to the PV in non-phagocytic cells, suggesting that GBPs do not localize to the PV during *Leishmania* infection (31). This data is in sharp contrast to localization studies performed on *T. gondii* infected bone marrow-derived monocytes and DCs (BM-MoDCs). During BM-MoDCs infection with *T. gondii*, GBP2 forms a tight ring around the PV containing *T. gondii* and iNOS is closely associated with GBP2^+^ PVs. These findings indicate that while GBPs may coordinate iNOS expression in our model, this coordination is not directly linked to GBP localization like during *T. gondii* infection. Importantly, the potential role of GBPs in parasite control uncoupled from their localization is a novel aspect of our research, as most studies have focused on the localization of GBPs to the PV or pathogen membrane.

## Conclusions

Ultimately, this work uncovers a critical host macrophage intrinsic mechanism in the control of *L. major* parasites. Current anti-leishmanial therapies, such as miltefosine, promote the production of IFN-γ, which subsequently induces ROS and NO production (65). However, the extent to which the therapeutic effects of these treatments are mediated through the activation of GBPs remains unclear. Future studies should focus on characterizing GBP responses during anti-leishmanial treatment to elucidate their potential role in enhancing parasite control. Targeting GBPs may hold therapeutic promise by promoting the expansion of iNOS^+^ macrophages, thereby improving the host’s ability to combat *L. major* and potentially other *Leishmania* species.

## Acknowledgments

This work was supported by the Center for Microbial Pathogenesis and Host Inflammatory Responses (funded by NIH NIGMS Centers of Biomedical Research Excellence Grant P20-GM103625). This publication was also supported in part by funds provided by the National Center for Advancing Translational Sciences of the NIH under awards TL1 TR003109 and UL1 TR003107 for the Systems Pharmacology and Therapeutics (SPaT) NIH T32 training grant GM106999 to Lucy Fry. The content is solely the responsibility of the authors and does not necessarily represent the official views of the NIH. The funders had no role in study design, data analysis, decision to publish or preparation of the manuscript.

